# Pharmaco-resistant temporal lobe epilepsy gradually perturbs the cortex-wide excitation-inhibition balance

**DOI:** 10.1101/2024.04.22.590555

**Authors:** Ke Xie, Jessica Royer, Raul Rodriguez-Cruces, Linda Horwood, Alexander Ngo, Thaera Arafat, Hans Auer, Ella Sahlas, Judy Chen, Yigu Zhou, Sofie L. Valk, Seok-Jun Hong, Birgit Frauscher, Raluca Pana, Andrea Bernasconi, Neda Bernasconi, Luis Concha, Boris Bernhardt

**Affiliations:** McConnell Brain Imaging Centre, Montreal Neurological Institute and Hospital, McGill University, Montreal, QC, Canada; Otto Hahn Research Group for Cognitive Neurogenetics, Max Planck Institute for Human Cognitive and Brain Sciences, Leipzig, Germany; Institute of Neurosciences and Medicine (INM-7), Research Centre Jülich, Jülich, Germany; Institute of Systems Neuroscience, Heinrich Heine University Düsseldorf, Düsseldorf, Germany; Center for Neuroscience Imaging Research, Institute for Basic Science, Sungkyunkwan University, Suwon, South Korea; Department of Biomedical Engineering, Sungkyunkwan University, Suwon, South Korea; Center for the Developing Brain, Child Mind Institute, New York City, NY, USA; Department of Neurology and Department of Biomedical Engineering, Duke University, Durham, NC, USA; Montreal Neurological Institute and Hospital, McGill University, Montreal, QC, Canada; Institute of Neurobiology, Universidad Nacional Autónoma de Mexico, Queretaro, Mexico

## Abstract

Excitation-inhibition (E/I) imbalance is theorized as a key mechanism in the pathophysiology of epilepsy, with a mounting body of previous research focusing on elucidating its cellular manifestations. However, there are limited studies into E/I imbalance at macroscale and its microcircuit-level mechanisms and clinical associations. In our current work, we computed the Hurst exponent—a previously validated index of the E/I ratio—from resting-state fMRI time series, and simulated microcircuit parameters using biophysical computational models. We found a broad reduction in the Hurst exponent in pharmaco-resistant temporal lobe epilepsy (TLE), indicative of a shift towards more excitable network dynamics. Connectome decoders pointed to temporolimbic and frontocentral areas as plausible network epicenters of E/I imbalance. Computational simulations further revealed that enhancing cortical excitability in patients likely reflected atypical increases in recurrent connection strength of local neuronal ensembles. Moreover, mixed cross-sectional and longitudinal analyses revealed heightened E/I elevation in patients with longer disease duration, more frequent electroclinical seizures and inter-ictal epileptic spikes, and worse cognitive functioning. Replicated in an independent dataset, our work provides compelling *in-vivo* evidence of a macroscale shift in E/I balance in TLE patients that undergoes progressive changes and underpins cognitive impairments, potentially informing treatment strategies targeting E/I mechanisms.

## Introduction

The balance between excitatory and inhibitory (E/I) signaling is a key principle of neuronal dynamics and cortical circuit function,^1, 2^ and plays a crucial role in typical neurodevelopment and the emergence of large-scale network coordination.^3, 4^ Conversely, imbalances in E/I have been implicated in numerous neurodevelopmental conditions.^5–8^ In particular, epilepsy constitutes a prototype condition of E/I imbalance. Here, E/I imbalances across different brain systems result in spontaneous seizures as well as inter-ictal epileptic phenomena, and can also impart cognitive and psychosocial consequences in patients.^9–11^ Although the pathophysiological mechanisms by which structural and functional brain changes in epilepsy^12–14^ cause epileptogenic events remain incompletely understood, E/I imbalance emerging from localized as well as distributed networks likely acts as a driver.^15, 16^ In temporal lobe epilepsy (TLE), the most common pharmaco-resistant focal epilepsy in adults, E/I imbalance is thought to originate primarily from temporolimbic circuits.^17, 18^ However, insights into the role of E/I dysfunction in TLE stem mainly from experimental studies in animal models and *ex-vivo* human tissue samples. *In-vivo* investigations in living human brains remain scarce thus far because of the limited availability of robust E/I biomarkers that are non-invasive, applicable in humans, and measurable on a large scale.

Functional magnetic resonance imaging (fMRI) provides a unique window into localized and macroscale functional properties in the living human brain.^19, 20^ More recently, advances in fMRI acquisition, processing, and signal modeling have permitted to approximate the E/I ratio with high spatial specificity and biophysical plausibility.^5, 21^ One recent study has proposed the Hurst exponent, a statistical descriptor of the spectral properties of neural time series, as an *in-vivo* neuroimaging biomarker that can index the synaptic E/I ratio.^5^ The Hurst exponent measures a signal’s fractal property to quantify temporal autocorrelation processes that occur within the signal itself.^22, 23^ *In-silico* modeling studies have confirmed a link to the E/I ratio, suggesting that increases in excitation within recurrent networks lead to flattening of 1/f slopes and reductions in the Hurst exponent.^5, 21^ Moreover, chemogenetic studies have shown that enhancing the excitability of excitatory neuronal populations in medial prefrontal regions reduces the Hurst exponent.^5^ Taken together, given its strong relevance to synaptic E/I mechanisms, the Hurst exponent can be deployed on a large scale for *in-vivo* investigation of E/I function in humans with TLE. The brain, and particularly the cortex, is organized hierarchically:^24^ cortical neurons assemble locally into microscale circuits that interconnect to form nodes, which in turn assemble to constitute macroscale networks. To gain a deeper understanding of the complex interplay between brain activity (*i.e.*, fMRI) and pathophysiological processes, models can provide more biologically plausible insights by incorporating heterogeneity of brain dynamics based on empirical data.^25^ In particular, neural mass models (NMMs) governed by anatomical and functional properties can robustly simulate interregional intrinsic functional connectivity from structural connectivity in healthy individuals.^26^ Moreover, model inversion techniques allow for the estimation of region-specific microcircuit parameters, such as recurrent connection strength and external subcortical input. NMMs-based whole-brain modeling has proven especially successful in simulating pathological perturbations of excitatory and inhibitory neurons in diseases like autism,^27^ Alzheimer’s disease,^28, 29^ and epilepsy.^30, 31^ Notably, patients with focal and generalized epilepsies have been shown to present with divergent patterns of cortical recurrent connection strength and subcortical inputs.^30, 31^ However, there remain critical gaps in the understanding of the neural mechanisms of cortical E/I imbalance in TLE and their potential clinical relevance.

Seizures have been established to increase markers of excitability, such as glutamate.^32–34^ Furthermore, TLE has consistently been associated with disruptions in glutamatergic and GABAergic circuits,^35, 36^ potentially contributing to the genesis or maintenance of seizure activity. Excessive metabolic activation resulting from disrupted balance in these systems may, in turn, promote excitotoxicity, epileptogenicity, and cell death.^37^ Ultimately, this process may lead to rapid seizure spread and an extension of the epileptogenic networks, affecting both seizure-generating and contralateral target regions.^38–41^ Despite a growing body of findings of progressive cortical atrophy in intractable TLE,^42–44^ it remains unclear whether recurrent seizures (or disease severity) in these patients are associated with progressive dysfunction of E/I balance. Examining whether changes in the Hurst exponent are more prominent in patients with longer duration of illness will shed new light on the progression of E/I imbalance. In this context, longitudinal designs provide an opportunity to infer causality between seizures and E/I imbalance.^42, 43^ Moreover, such designs can control for aging effects and inter-subject variability, thereby increasing sensitivity to detect subtle changes. In addition to experiencing seizures, TLE patients are also affected by cognitive, psychological, and social impairment. Up to 80% of patients demonstrate impairments in at least one cognitive domain—most frequently memory, executive, and language function.^45–48^ In a subset of patients, these impairments have been shown to be progressive in nature.^49–51^ Despite the high prevalence of cognitive dysfunction in TLE, there is significant variability in the severity of impairments observed across patients. For example, patients with generalized cognitive impairment demonstrate widespread cortical thinning and diffuse white matter compromise,^46, 51, 52^ whereas those with intact cognitive profiles have minimal structural abnormalities. Emerging evidence suggests that cognitive impairment in TLE is also determined by damage to functional connectivity within the medial temporal lobe.^14, 53^ To date, however, no studies have explored the extent to which E/I changes (*i.e.*, the Hurst exponent) can predict cognitive impairments in TLE.

In this study, we profiled cortical E/I imbalance patterns in pharmaco-resistant TLE patients and determined their associations with microcircuit perturbations and clinical presentations. We derived the region-wise Hurst exponent from resting-state fMRI time series as an E/I ratio proxy and compared this metric between TLE patients and matched healthy controls. Subsequently, we employed biophysical computational simulations to elucidate microcircuit-level mechanisms (recurrent connection strength and external input) underlying macroscale E/I imbalance across the brain. Additionally, we explored associations between E/I ratio changes and brain perfusion alterations as well as electroclinical parameters. Finally, to demonstrate clinical relevance, we assessed the progression of E/I imbalance in our patient cohort, and its relation to clinical scores of disease severity and cognitive function using both cross-sectional and longitudinal designs. The reproducibility of our findings was verified in an independent validation dataset.

## Results

We analyzed 2 independent datasets with multimodal MRI data (*i.e.*, structural, diffusion, and restingstate fMRI). The discovery dataset *MICA-MICs*, collected at Montreal Neurological Institute-Hospital,^54^ included 80 participants (40 healthy controls, 40 pharmaco-resistant TLE). The replication dataset *EpiC* included 60 participants (30 healthy controls, 30 pharmaco-resistant TLE) from Universidad Nacional Autónoma de México.^48, 51^ TLE diagnosis was determined according to the classification of the ILAE.^55^ Details of subject inclusion criteria are provided in **Methods**. Site-specific demographical and clinical information are shown in **Table 1**. All participants were aged between 18 and 63 years, with no significant group differences observed for age or sex.

**Table 1.** Demographic and clinical information.

|  | Discovery Dataset (*MICA-MICs*) |  |  | Replication Dataset (*EpiC*) |  |  |
| --- | --- | --- | --- | --- | --- | --- |
|  | Controls | TLE | *P*-value | Controls | TLE | *P*-value |
| Number | 40 | 40 | - | 30 | 30 | - |
| Age | 34.25 ± 3.98<br>(28 - 44) | 35.80 ± 11.04<br>(18 - 63) | 0.406<sup>a</sup> | 31.83 ± 11.35<br>(18 - 57) | 30.87 ± 11.46<br>(18 - 58) | 0.744<sup>a</sup> |
| Sex (M/F) | 19/21 | 17/23 | 0.653<sup>b</sup> | 11/19 | 10/20 | 0.787<sup>b</sup> |
| Focus (L/R) | - | 27/13 | - | - | 18/12 | - |
| Age at seizure onset (years) | - | 21.80 ± 11.24<br>(6 - 60) | - | - | 14.96 ± 10.51<br>(0.7 - 40) | - |
| Duration of epilepsy (years) | - | 14.00 ± 11.27<br>(1 - 45) | - | - | 15.58 ± 13.14<br>(1 - 49) | - |
| AEDs | - | 2.28 ± 0.85<br>(1 - 5) | - | - | 1.62 ± 0.64<br>(1 - 3)<sup>d</sup> | - |
| Surgery (Engel I) | - | 16 (11)<sup>c</sup> | - | - | - | - |

Age, age at seizure onset, and duration of epilepsy are presented as mean ± SD years. <sup>a</sup> Two-sample *t*-test. <sup>b</sup> Chi-square test. <sup>c</sup> Engel I: seizure-free, *i.e.*, Class I postsurgical outcome in Engel's classification. <sup>d</sup> Information available in 26 TLE patients. TLE = temporal lobe epilepsy; M = male; F = female; L = left; R = right; AEDs = anti-epileptic drugs.

### Hurst Exponent Reductions in TLE

We calculated the region-wise Hurst exponent value, a metric that is mathematically related to the 1/f exponent of neural signals^21, 56^ and used as an index of the E/I ratio. As previously described,^5, 57^ resting-state fMRI time series were modeled as multivariate fractionally integrated processes, and the Hurst exponent was estimated via the univariate maximum likelihood method and discrete wavelet transform.^5^ As such, an elevated E/I ratio would manifest in a lower Hurst exponent value. In healthy control and TLE groups reported here, interregional variations in the Hurst exponent values exhibited hierarchical gradients, with the highest values observed in the sensory regions, intermediate values in the association regions, and lowest values in the paralimbic regions (**Fig. 1a, 1c**). This pattern of sensory-fugal distinction was confirmed by a significant spatial correlation with the principal axis of cytoarchitectural differentiation (*rho* = −0.41, *P*_spin_ = 0.044; **Supplemental Fig. 1a**), previously established through analysis of myelin-sensitive MRI.^58^ To further contextualize our regional pattern of the Hurst exponent, we correlated it with morphometric and molecular markers of phylogenetic cortical differentiation.^59^ We found that at the surface level, the Hurst exponent positively correlated with intracortical myelination (*rho* = 0.37, *P*_spin_ = 0.066) and the gene expression gradient (*rho* = 0.60, *P*_spin_ = 0.002),^60^ while negatively correlating with cortical thickness (*rho* = −0.51, *P*_spin_ = 0.007) and the neurotransmitter receptor gradient (*rho* = −0.46, *P*_spin_ = 0.009;^61^ **Supplemental Fig. 1b**).

**Fig. 1.**
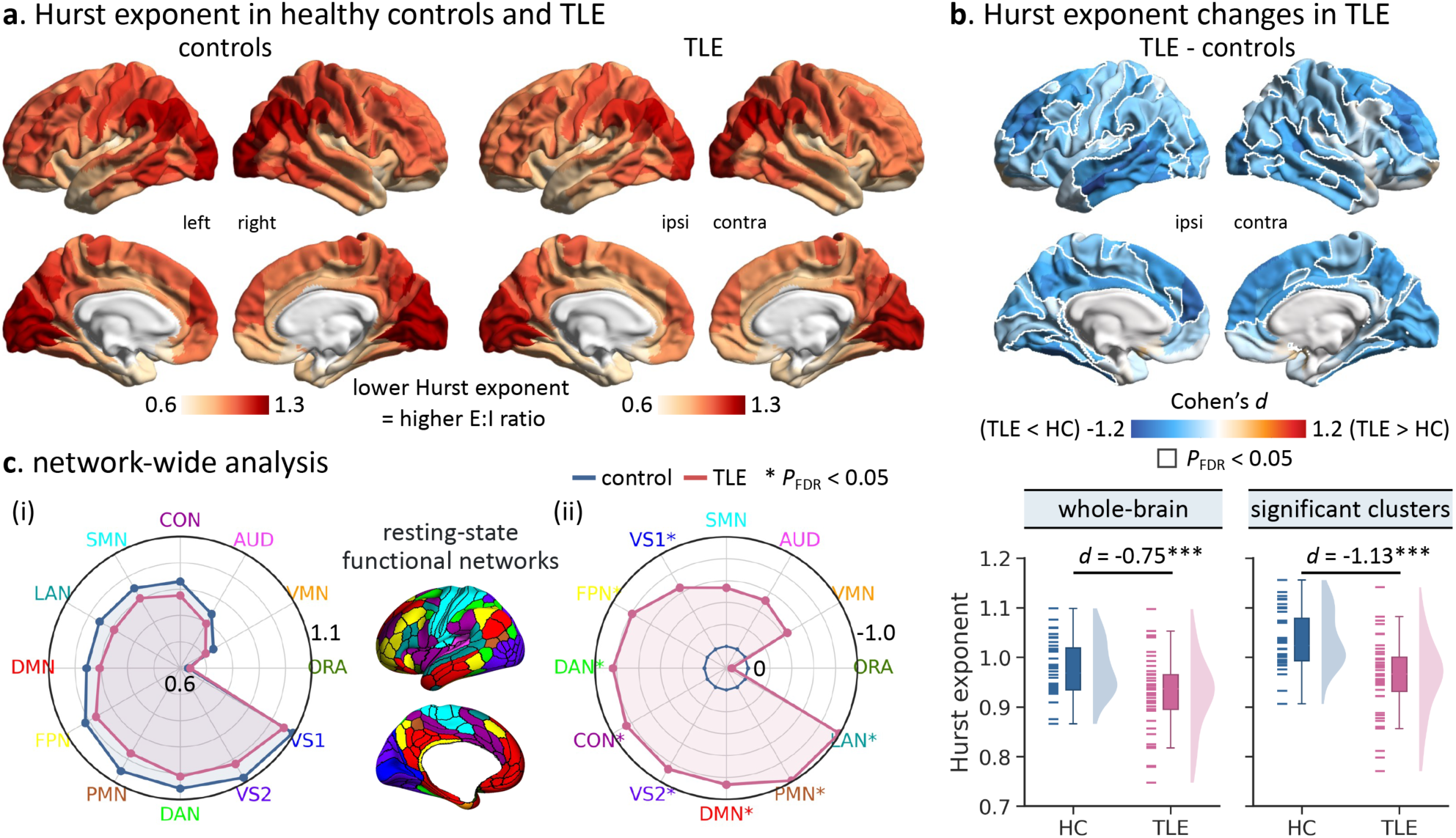
Hurst exponent reductions in TLE. **(a)** Mean regional patterns of the Hurst exponent of resting-state fMRI time series in healthy controls and TLE patients: the lower the Hurst exponent, the higher the excitation/inhibition ratio.^5,57^ **(b)** Top: statistical map of TLE-control difference in regional Hurst exponent, effect size as Cohen’s *d*. Significant regions, corrected for multiple comparisons using the false discovery rate procedure (*P*_FDR_ < 0.05), are surrounded by solid white outlines. Bottom: participant-specific average Hurst exponent across the whole brain or statistically significant regions, respectively. **(c)** (i) Distribution of the average Hurst exponent in 12 large-scale functional networks in healthy control and TLE groups, respectively. (ii) Distribution of TLE-control differences in the Hurst exponent with respect to each network (*P*_FDR_ < 0.05).^62^ *** *P* < 0.001; HC = healthy controls; TLE = temporal lobe epilepsy; ipsi = ipsilateral; contra = contralateral; VS1/VS2 = primary/secondary visual network; AUD = auditory network; SMN = somatomotor network; CON = cingulo-opercular network; DAN = dorsal attention network; LAN = language network; FPN = fronto-parietal network; DMN = default mode network; PMN/VMN = posterior/ventral multimodal network; ORA = orbito-affective network.

In comparison to healthy controls, TLE patients exhibited marked reductions in the Hurst exponent both at global and local levels. Specifically, TLE patients had a significantly lower grand average Hurst exponent across the entire brain relative to healthy controls (Cohen’s *d* = −0.75, *P* < 0.001; **Fig. 1b**). Surface-based analysis showed significantly a lower Hurst exponent in 156 out of 360 brain regions in TLE compared to healthy controls following correction for multiple comparisons at a false discovery rate of *P*_FDR_ < 0.05 (**Fig. 1b**). These mostly affected the lateral inferior, middle and superior temporal lobes, the dorsolateral and dorsomedial prefrontal cortices, the fusiform gyrus, the precuneus, and the occipital cortex bilaterally, together with the ipsilateral postcentral gyrus, with effect sizes ranging from medium to large (*d* = −0.46 - −1.22, mean ± SD = −0.64 ± 0.07). These findings were verified in a subgroup TLE patients with histologically confirmed mesiotemporal sclerosis and post-surgical seizure freedom at a 1-year follow-up (*i.e.*, Engel I, *n* = 11; whole-brain/significant clusters: *d* = −1.56/-1.94, *P* < 0.001; **Supplemental Fig. 2**). Finally, when summarizing the cortex-wide findings with respect to 12 intrinsic functional communities,^62^ pronounced between-group effects were seen in the transmodal association system, such as the default mode, frontoparietal, and dorsal attention networks, as well as the unimodal visual system (*P*_FDR_ < 0.05; **Fig. 1c**). To ensure that our results were not related to spurious features, we assessed the degree of head motion of each individual during resting-state fMRI scan based on framewise displacement (FD).^63^ Notably, between-group differences in the global average (*d* = −0.47, *P* = 0.020) and local value of the Hurst exponent (mean ± SD *d* = −0.58 ± 0.07; **Supplemental Fig. 3**) were robust when additionally controlling for individual-wise mean FD, suggesting no marked influence of head motion.

### Microcircuit Parameter Changes in TLE

Next, we cross-validated our findings and explored circuit mechanisms underlying disruptions of E/I balance in TLE via a parametric mean-field model (pMFM).^26^ In the biophysically-based model, local neuronal dynamics were simulated through a set of simplified nonlinear stochastic differential equations (see **Methods**) by linking ensembles of local neuronal masses with diffusion-derived structural connectivity.^64^ The pMFM iteratively tuned its parameters (*i.e.*, recurrent connection strength *w*, external input current *I*, noise *σ*, and a constant *G*) to simulate neural signals that were maximally similar to the empirical data. Adopting a recent framework,^26^ *w*, *I*, and *σ* varied across brain regions and were parameterized by a linear combination of local structural (*i.e.*, intracortical myelination) and functional (*i.e.*, resting-state functional connectivity gradient) properties (**Fig. 2a**). More specifically, for each group (healthy control or TLE), the 40 participants were randomly subdivided into the training (*n* = 15), validation (*n* = 15), and test (*n* = 10) sets. Group-averaged structural connectivity, static functional connectivity (FC), and time-varying functional connectivity dynamics (FCD) were computed separately for each set. Next, the 250 candidate parameter sets (*w*, *I*, *σ*, and *G*) were generated from the training set of each group using the CMA-ES algorithm and evaluated in the validation set.^65^ The top 10 parameter sets from the validation set were then evaluated in the test set. To ensure stability, the split of participants into training, validation, and test sets was repeated five times. Finally, the pMFM parameters based on the best fit (with the lowest cost between simulated and empirical FC and FCD matrices; see **Methods**) from the test set were averaged across these five splits, yielding the representative set of parameters for each group.

**Fig. 2.**
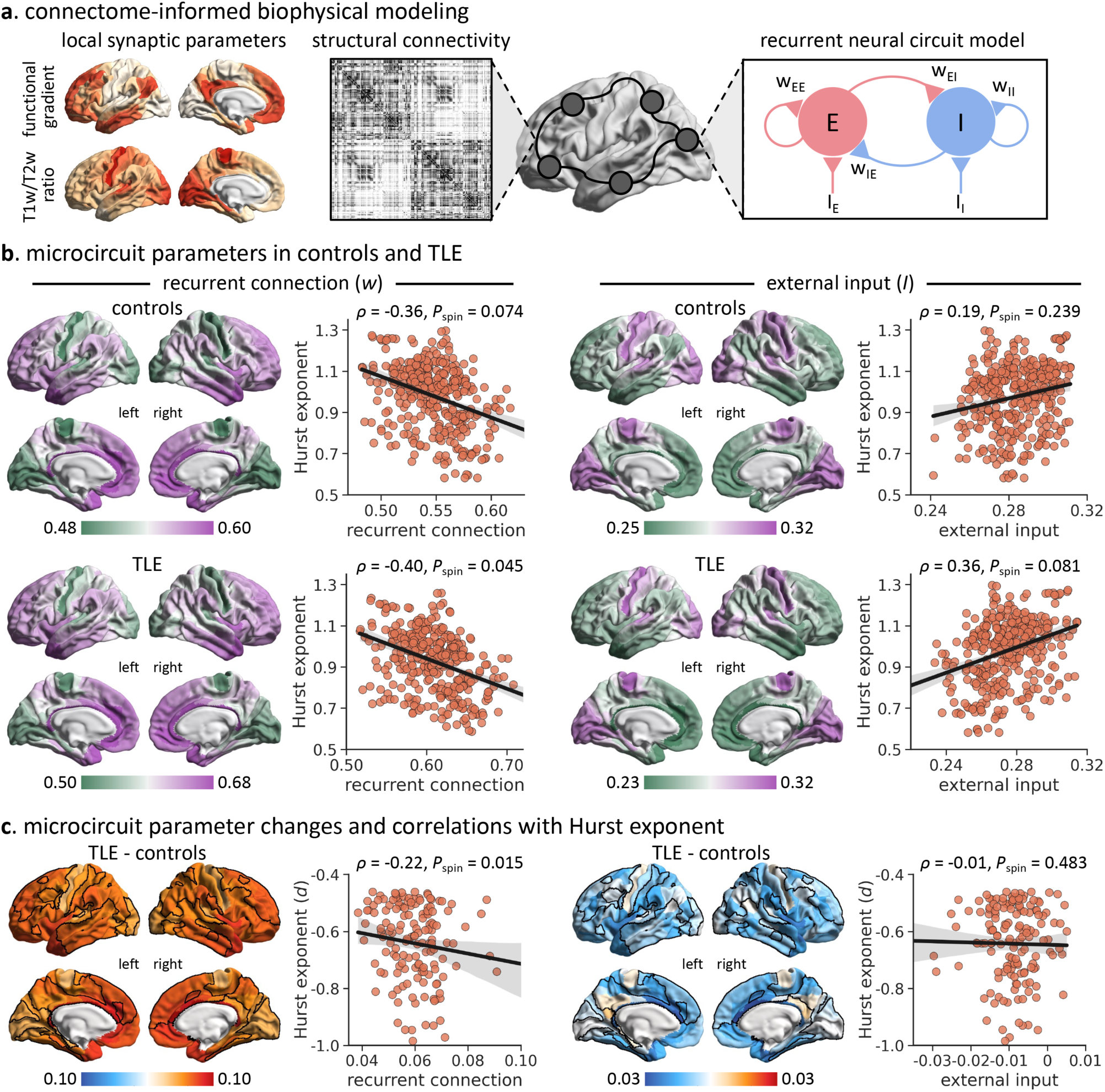
Microcircuit parameter differences in TLE and associations with Hurst exponent changes. **(a)** Schematic of the parametric mean-file model (pMFM) to estimate region-specific microcircuit parameters (*i.e*., recurrent connection strength *w* and subcortical input *I*) from structural connectivity.^26^ The pMFM is parameterized by a linear combination of resting-state functional connectivity gradient and T1w/T2w ratio. *E* and *I* indicate the excitatory and inhibitory neuronal populations, respectively. *W*_EI_ indicates the strength of connection from the excitatory population to the inhibitory population, and so forth. **(b)** Regional microcircuit parameters of healthy controls (top) and TLE patients (bottom) and associations with the Hurst exponent at the surface level. **(c)** Regional changes in microcircuit parameters (TLE-control) and relations to the effect sizes of Hurst exponent alterations (*i.e*., Cohen’s *d* scores), constrained to brain regions showing significant between-group differences in Fig. 1b (solid black outlines). The statistical significance of spatial correlation between spatial maps (*i.e*., *P*_spin_) is determined via spin permutation tests (5,000 iterations).^66, 67^

In agreement with prior work,^26^ in both cohorts, recurrent connection strength (*w*) gradually increased from primary sensory/motor regions to high-order association regions across the neocortex, reaching its highest value in the default mode network. On the other hand, external input (*I*) smoothly decreased along the unimodal-transmodal hierarchy (**Fig 2b**). By correlating regional variations in microcircuit parameters with the Hurst exponent, we observed that recurrent connection strength (*w*) appeared to be stronger in brain areas with a lower Hurst exponent (*i.e*., higher E/I ratio) (healthy controls: *rho* = −0.36, *P*_spin_ = 0.074; TLE: *rho* = −0.40, *P*_spin_ = 0.045; **Fig 2b**). On the other hand, regions with higher local external input (*I*) had a relatively higher Hurst exponent (*i.e*., lower E/I ratio) (healthy controls: *rho* = 0.19, *P*_spin_ = 0.239; TLE: *rho* = 0.36, *P*_spin_ = 0.081; **Fig 2b**). Moreover, compared to healthy controls, TLE patients showed increases in recurrent connection strength (*w*) as well as decreases in external input (*I*). TLE-related reductions in the Hurst exponent were enriched in areas with the greatest effects of increasing recurrent connection strength (*rho* = −0.22, *P*_spin_ = 0.015). By contrast, no significant correlation was found between the degree of the Hurst exponent alterations and regional changes in external input (*rho* = −0.01, *P*_spin_ = 0.483; **Fig 2c**).

### Network-Level Effects of Hurst Exponent Reductions

Local vulnerability interacts with brain network architecture to shape disease pathology and spread.^68–71^ Here, we assessed the extent to which TLE-related changes in the Hurst exponent exhibited network effects. For each node, we computed the mean Hurst exponent change (*i.e.*, mean Cohen’s *d*) of its connected neighbours, weighted by streamline density estimated using diffusion MRI and functional connectivity strength estimated using resting-state fMRI (**Fig. 3a**). To ensure that connectivity estimates reflect the typical connectomes prior to disease onset and deafferentation, we estimated group-level structural and functional connectivity matrices in a sample of 100 unrelated healthy young adults from the Human Connectome Project (S900-HCP).^72^ We observed a positive association between the change in the Hurst exponent of a node and the mean change of its structurally connected neighbours (*rho* = 0.54, *P*_spin_ < 0.001; **Fig. 3a**), above and beyond the effect of spatial autocorrelation. Similarly, we found a comparable association between the change in the Hurst exponent of a node and the mean change of its functionally connected neighbours (*rho*= 0.47, *P*_spin_ < 0.001; **Fig. 3a**). That is, pathology in a brain region is closely correlated with greater exposure to pathology in anatomically and/or functionally connected regions.

**Fig. 3.**
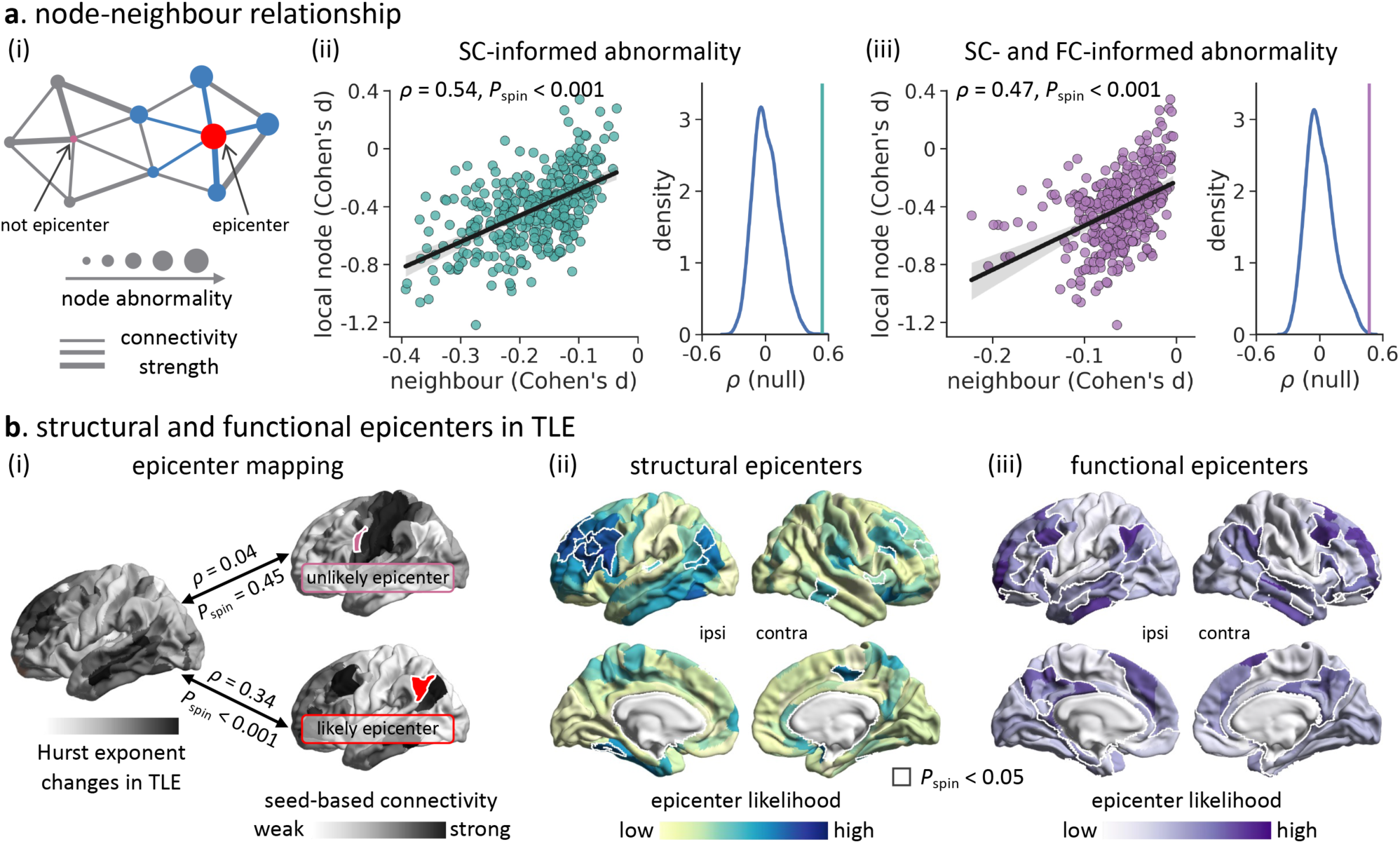
Network-based spreading of Hurst exponent reduction. **(a)** (i) Schematic of structural (SC) or functional (FC) connectivity informing TLE-related Hurst exponent changes. (ii-iii) Correlation between node abnormality and SC-/FC-informed mean neighbour abnormality. The histogram indicates the null distribution of correlation coefficients generated based on 5,000 spin tests, and the actual *r* value is represented by a green/purple bar. **(b)** Epicenters of TLE-related Hurst exponent changes. (i) A node whose SC or FC pattern across the entire cortex strongly correlated with TLE-related Hurst exponent change map (see Fig. 1b) is considered a likely “epicenter”. Epicenter likelihood was defined as the correlation coefficient between the two maps. (ii-iii) SC- and FC-informed epicenter likelihood maps of TLE-related Hurst exponent reductions, where the most likely epicenters, assessed using spin permutation tests (5,000 iterations, and *P*_spin_ < 0.05), are surrounded by solid white outlines. ipsi = ipsilateral; contra = contralateral.

Having observed that network architecture reflects TLE-related Hurst exponent change, we then examined which brain regions likely act as putative disease epicenters. As previously introduced,^44, 71^ we defined an epicenter as a node wherein structural and functional connectivity profiles spatially resembled the whole-brain pattern of TLE-related Hurst exponent changes (**Fig. 3b**). This measure identifies “disorder hubs”—regions that are both vulnerable to disorder-specific changes but also embedded in a highly atypical network cluster. Nodes were ranked in descending order based on their correlation coefficients. Empirical epicenter likelihood rankings were compared with rankings estimated from spatial autocorrelation-preserving null models (5,000 iterations). Several brain areas emerged as potential epicenters, including the bilateral dorsolateral prefrontal, medial and inferior temporal lobe, the precuneus, and the superior parietal cortex (*P*_spin_ < 0.05; **Fig. 3b**).

### Associations with Clinical and Cognitive Variables

Associations between changes in Hurst exponent and clinical characteristics of disease severity were assessed in TLE patients. A longer disease duration was negatively correlated with the Hurst exponent (whole-brain: *t* = −1.91, *P* = 0.016; significant clusters: *t* = −2.02, *P* = 0.013; **Fig 4a**), indicating relatively lower Hurst exponent in patients with long-standing TLE. There were also negative correlations between the Hurst exponent and the number of electroclinical seizures captured during hospitalization (*r* = −0.38, *P* = 0.009; *r* = −0.36, *P* = 0.013; **Supplemental Fig. 4**), such that more frequent epileptic seizures were associated with lower Hurst exponent. Similarly, we found significantly lower Hurst exponent values in TLE patients with prevalent interictal epileptic discharges (IEDs) than those with rare IEDs (*d* = −0.70, *P* = 0.029; *d* = −0.67, *P* = 0.034; **Supplemental Fig. 5**). No significant correlations were found between the Hurst exponent and age at seizure onset or the number of antiepileptic drugs (*P* > 0.320). To further understand the neurophysiological substrate of TLE-related changes in the E/I ratio, we analyzed cerebral blood flow measured by pseudo-continuous arterial spin labeling (ASL) MRI in patients with TLE. We found significantly lower cortical perfusion (or hypoperfusion) in regions (*rho* = 0.41, *P*_spin_ < 0.001), or patients with lower Hurst exponent values (whole-brain: *r* = 0.40, *P* < 0.001; significant clusters: *r* = 0.43, *P* < 0.001; **Supplemental Fig. 6**).

**Fig. 4.**
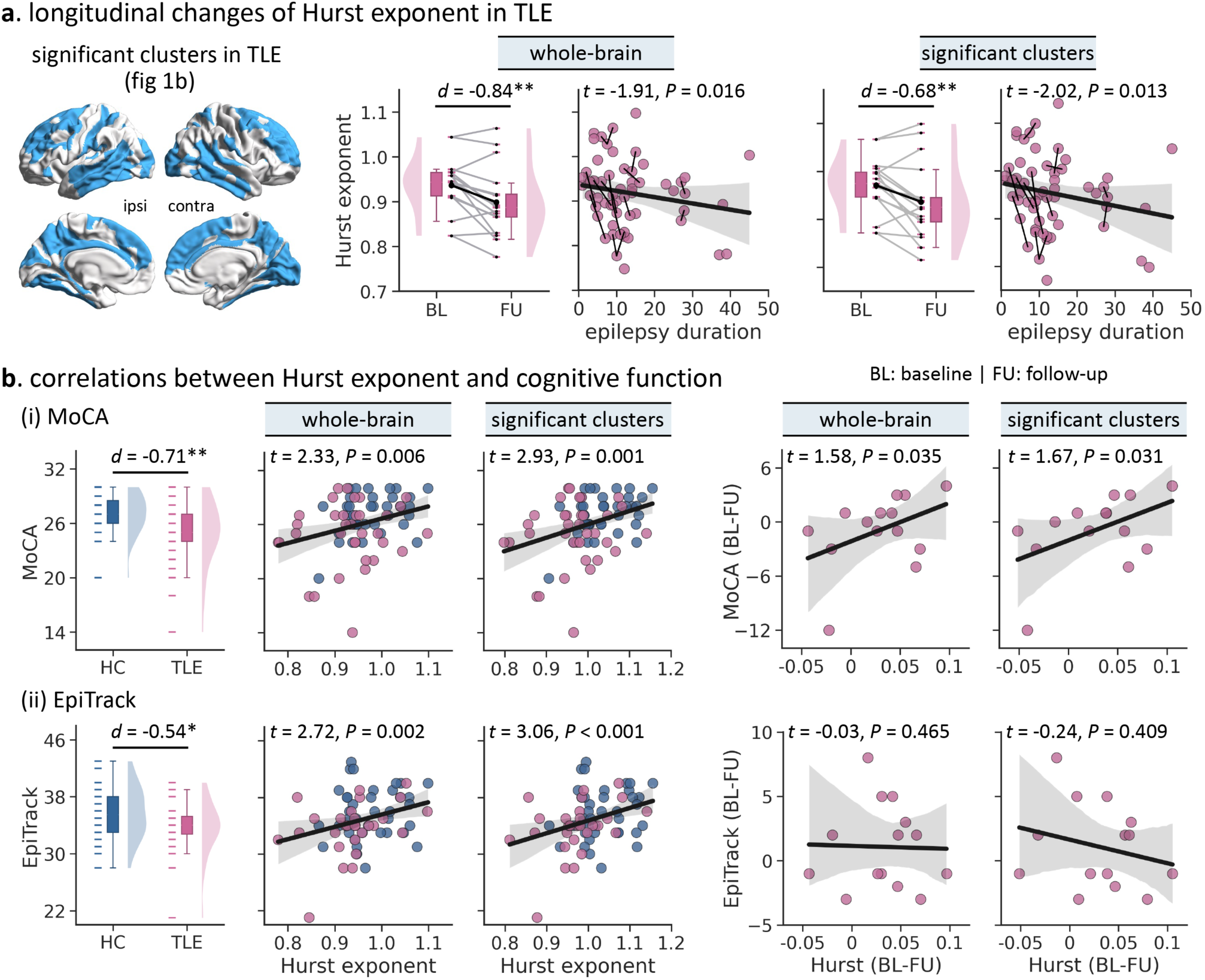
Associations of the Hurst exponent with clinical characteristics and behavioral assessments. **(a)** Error bar plots: longitudinal changes in the Hurst exponent in TLE patients. Spaghetti plots: associations between epilepsy duration and the Hurst exponent across both baseline and follow-up time points. **(b)** Left: MoCA and EpiTrack scores in TLE patients and healthy controls at baseline. Middle: associations between the Hurst exponent with MoCA and EpiTrack scores at baseline. Right: associations between longitudinal changes in the Hurst exponent, and MoCA and EpiTrack scores in TLE patients. * *P* < 0.05; ** *P* < 0.01. HC = healthy controls; TLE = temporal lobe epilepsy; MoCA = Montreal Cognitive Assessment; BL = baseline; FU = follow-up.

We then explored the associations between the Hurst exponent and cognitive function in TLE patients both cross-sectionally and longitudinally. As previously reported,^73^ at baseline, TLE patients showed markedly poorer performance than healthy controls in general cognitive functioning as measured by the MoCA test^74^ (*d* = −0.71, *P* = 0.002), as well as in attention and executive functions as measured by the EpiTrack test^75^ (*d* = −0.54, *P* = 0.017; **Fig. 4b**). The Hurst exponent positively correlated with MoCA scores (whole-brain: *t* = 2.33, *P* = 0.006; significant clusters: *t* = 2.93, *P* = 0.001), as well as EpiTrack scores (*t* = 2.72, *P* = 0.002; *t* = 3.06, *P* < 0.001; **Fig. 4b**), indicating more marked cognitive impairment in patients with lower Hurst exponent values. Furthermore, by analyzing for whom longitudinal data was available, we found that the Hurst exponent progressively decreased over a mean interscan interval of 2 years (whole-brain: *d* = −0.84, *P* = 0.002; significant clusters: *d* = −0.68, *P* = 0.006; **Fig. 4a**), and correlated with the progressive decline in MoCA scores (*t* = 1.58, *P* = 0.035; *t* = 1.67, *P* = 0.031; **Fig. 4b**).

### Replication Analysis

Hurst exponent reductions in TLE were replicable in an independent replication dataset (*i.e.*, EpiC) consisting of 30 pharmaco-resistant TLE patients and 30 healthy controls. Specifically, in the *EpiC* dataset, the spatial pattern of the Hurst exponent in each group closely resembled that observed in the *MICA-MICs* dataset, gradually decreasing along the sensory-fugal axis (*rho* > 0.89, *P*_spin_ < 0.001; **Fig. 5a**). Comparing cohorts in the *EpiC* dataset, TLE patients also exhibited significantly decreased Hurst exponent globally (*d* = −0.61, *P* = 0.011), as well as locally in significant regions identified from the discovery dataset (*d* = −0.60, *P* = 0.012; **Fig. 5b**). Region-wide group differences in the Hurst exponent were also spatially correlated between the two datasets at the surface level (*rho* = 0.20, *P*_spin_ = 0.010). Furthermore, we observed a noticeable trend toward progressive decreases in the Hurst exponent in TLE patients over a mean interscan period of 2.5 years (whole-brain: *d* = −0.35, *P* = 0.105; significant clusters: *d* = −0.38, *P* = 0.087). A longer epilepsy duration was associated with a greater extent of decreases in the Hurst exponent (*t* = −3.33, *P* = 0.001; *t* = −3.28, *P* = 0.002; **Fig. 5c**). Finally, we replicated the finding of the cognitive relevance of an altered Hurst exponent, with evidence of marked memory deficits in TLE compared to healthy controls (PC1 score: *d* = −0.91, *P* < 0.001), as well as lower Hurst exponent values in TLE with poorer memory performance (*t* = 2.06, *P* = 0.011; *t* = 2.05, *P* = 0.012; **Fig. 5d**).

**Fig. 5.**
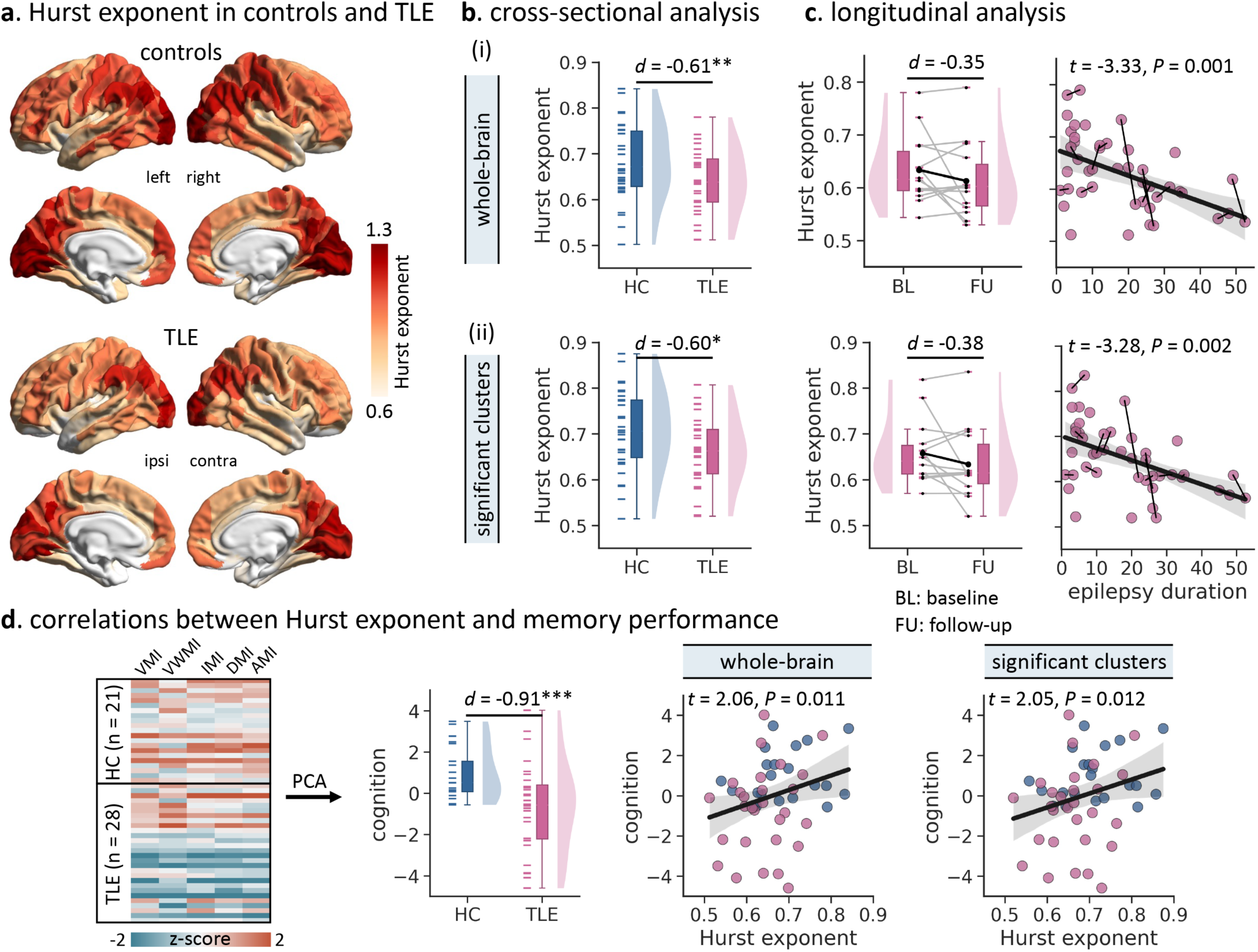
Replication analysis of the Hurst exponent changes in *EpiC* dataset. **(a)** Group-averaged Hurst exponent in healthy control and TLE groups. **(b)** TLE-control differences inthe average Hurst exponent across the entire brain (i) and significant clusters (ii) identified from the discovery dataset (see Fig. 1b). **(c)** Error bar plots: longitudinal changes in the Hurst exponent in TLE patients. Spaghetti plots: associations between epilepsy duration and the Hurst exponent across baseline and follow-up time points. (**d**) Relationships between individual memory performance and the Hurst exponent at baseline. Error bar plots: between-group differences in the overall memory performance (*i.e.,* PC1 loading score) determined through a principal component analysis on various memory tests. Scatter plots: individual Hurst exponent positively correlated with the PC1 loading score after controlling for age and sex. * *P* < 0.050; ** *P* < 0.010; *** *P* < 0.001. HC = healthy controls; TLE = temporal lobe epilepsy; BL = baseline; FU = follow-up; AMI = auditory memory; DMI = delayed memory; IMI = immediate memory; VMI = visual memory; VWMI = visual working memory.

## Discussion

In this work, we mapped cortical E/I imbalance in patients with pharmaco-resistant TLE and assessed its relationships with microcircuit-level dysfunction and cognitive impairments. We found significant decreases in the Hurst exponent in TLE compared to controls in cortical networks that extended beyond the temporolimbic cortex to frontoparietal and occipital regions, suggesting a shift in E/I balance toward large-scale network excitation. Leveraging whole-brain biophysical simulations, we demonstrated that enhancing cortical excitation in TLE reflected atypical increases in recurrent connection strength within the structurally governed functional connectome. Moreover, mixed cross-sectional and longitudinal analysis unveiled more marked Hurst exponent decreases in TLE with longer disease duration, more frequent electroclinical seizures and inter-ictal epileptic spikes, as well as poorer cognitive function. The Hurst exponent showed a progressive decrease at longitudinal follow-up, and correlated with the simultaneous longitudinal worsening of cognitive impairments in patients. Our findings were replicated in an independent dataset, suggesting generalizability. Taken together, our work provides robust *in-vivo* evidence supporting the existence of cortical E/I balance shifting toward excitation in pharmaco-resistant TLE. These findings enhance our understanding of the interplay between macroscale functional imbalance, microcircuit perturbation, and cognitive dysfunction, potentially informing new treatment strategies targeting E/I mechanisms.

Our study examined the Hurst exponent, an *in-vivo* neuroimaging marker of E/I balance, in TLE. Prior research with simplified models has indicated that the Hurst exponent in neural time series data closely reflects underlying changes in synaptic E/I ratio.^5^ In recurrent networks, where excitatory and inhibitory neuronal populations interact, the Hurst exponent decreases with increasing network excitability. Although net E/I effects are typically balanced in local circuits in healthy individuals, there are slight variations in the degree of balance across regions. Specifically, in our control group, the Hurst exponent values gradually decreased along the sensory-fugal hierarchy,^58^ with the highest values observed in the primary sensory regions with heavy myelination and laminar organization, and the lowest values in paralimbic regions. Increased levels of myelination have been reported to suppress the formation of new axonal tracts and synapses, thus potentially reducing spikes and yielding neural activity with less scale-free or critical properties.^1, 76, 77^ By contrast, lower myelination in frontal and limbic cortices allows for greater functional signal variability and neuronal remodeling at various timescales, facilitating the emergence of diverse functional dynamics.^58, 78^ Strikingly, we found significantly lower Hurst exponent levels in bilateral frontal, central, temporal, and occipital regions in TLE patients compared to controls, with more pronounced changes ipsilateral to the seizure focus. These findings indicate that the pathophysiological substrate of TLE is likely defined by a network of multiple interconnected regions,^71^ rather than solely dependent on the mesiotemporal lobe. Our results are also compatible with existing *in-vitro* imaging data in TLE demonstrating overall hyperexcitability of local networks, which may be fostered by atypical excitatory and inhibitory processes at the cellular level.^10^ Although the current study examined a relatively homogenous cohort of patients with electroclinical features of unilateral TLE, alterations in the Hurst exponent encompassed a bilateral territory. Previous electrophysiology studies have shown that seizures originating in the ipsilateral mesiotemporal region often propagate to the contralateral temporal lobe directly along commissural pathways or indirectly via other regions, such as the frontal lobes.^79^ Ultimately, cellular and synaptic alterations may occur in both seizure-generating ipsilateral regions and contralateral zones of propagation.^80^ In TLE, increased slow waves are characteristic features during the interictal period; higher incidences of slow waves are related to greater volume loss in mesiotemporal structures.^81^ This suggests a potential link between neuronal death, synaptic loss, and cortical hyperactivity, which warrants future validation using pathology data from patients undergoing surgery. Moreover, datadriven epicenter mapping revealed that regional changes in the Hurst exponent implicated both structurally and functionally connected neighbours, suggesting that network architecture serves as a scaffold for the spread of E/I imbalance in TLE. Interestingly, bilateral temporolimbic and frontoparietal regions (precuneus, superior parietal cortex) emerged as putative epicenters. These regions are generally considered densely inter-connected hubs that are thought to support the integration and broadcasting of signals across different subnetworks.^82^ Hubs are particularly susceptible to pathology, with mounting evidence showing changes in cortical morphology and connectivity patterns in TLE.^20, 71, 83^ Indeed, fMRI studies in TLE have previously identified reductions in long-range connections of distributed cortical networks, alongside increases in connectivity in temporolimbic circuits proximal to the seizure focus.^84^ We complement these findings by highlighting that the temporolimbic and frontoparietal cortices are particularly vulnerable to E/I disturbance and that they, by virtue of their network embedding, may increase disease exposure to connected regions.

Deviations from macroscale E/I balance may be related to dysfunctions of neural circuits. Our work quantified the extent of perturbations in cortical microcircuit function in TLE, by leveraging a wholebrain computational model (pMFM) that is biophysically grounded yet parsimonious.^26^ In recent work, pMFM has demonstrated the ability to predict functional connectivity from structural connectome with robust accuracy and low parametric complexity. In our work, interregional variations in recurrent connection strength and external input current followed sensory-association hierarchical gradients, and exhibited significant correlations with Hurst exponent maps at the surface level. This suggests a potential link between microcircuit dynamics and cortex-wide heterogeneity in E/I balance. Importantly, comparing model parameters between TLE patients and controls suggested a diffuse pattern of local microcircuit disruptions, particularly marked in default mode and frontoparietal systems. Directly correlating TLE-related Hurst exponent changes with microcircuit parameters revealed a unique association with increased recurrent connection strength, which indicates aberrant excitatory function within local ensembles of neuronal subpopulations. As such, excessive intrinsic neuronal excitability may underpin the increased E/I ratio in TLE patients. This is consistent with multiple lines of evidence indicating how deficits in excitatory function may affect macroscale brain dynamics.^30, 31^ Basic science experiments have identified aberrant glutamate transmission,^85, 86^ as well as inhibitory interneuron hypofunction as potential underlying causes of neuronal hyperactivity, favoring recurrent seizure activity in TLE.^35, 36, 80^ Future work is warranted to more precisely delineate the contributions from excitatory and inhibitory sub-population functions toward network hyperexcitability, particularly given that the current study only considered the recurrent interactions of two canonical cell types. Different classes of inhibitory interneurons exhibit diverse cellular and synaptic properties, microcircuit connectivity patterns, and neurophysiological responses. Another future direction is to incorporate distinct classes of inhibitory interneurons into circuit models, allowing for a more nuanced investigation of inhibitory dysfunction beyond the relatively coarse net effect of the E/I ratio. Overall, our findings from computational simulations help bridge a crucial gap between microcircuit dysfunctions and perturbed E/I balance at the macro scale in TLE.

The utility of a biomarker is often contingent upon its relationship with clinical measures and core symptoms. By leveraging the considerable range of disease duration in our patient cohort, we found significantly lower global as well as local Hurst exponent values in individuals with longer duration, more frequent electroclinical seizures, and more abundant inter-ictal epileptic spikes. This finding is compatible with earlier cross-sectional structural work on TLE with wider windows of disease duration or stages, which demonstrated a greater extent and broader distribution of cortical thinning in subgroups with longer duration compared with those with shorter duration of disease.^71, 87–89^ Current findings, together with previous evidence of cumulative metabolic changes,^44, 90^ indicate that TLE is likely a progressive neurological disorder. Our longitudinal substudy supports this notion, indicating progressive reductions in the Hurst exponent in TLE over the follow-up period. Recurrent seizures and progressive E/I dysfunction may contribute to epileptic discharges and secondary damage in other regions and, in turn, extend epileptogenic networks. The static and dynamic nature of pathology in intractable TLE, as shown here and in other studies, underscores the critical importance of early surgical intervention in operative candidates to prevent adverse brain reorganization. In patients for whom surgery is delayed nevertheless, serial scanning may allow for establishing biomarkers of clinical outcomes. In addition, the extent of Hurst exponent decreases correlated with deficits in high-order cognitive functions in TLE, both in our cross-sectional and longitudinal studies. Previous studies have explored potential links between structural and functional alterations in TLE and impairment across various cognitive domains, including memory, language, and executive control.^14, 45, 47, 53^ Reduced volumes in subregions of the prefrontal cortex, for example, have been related to poor executive functioning^91^ and impaired memory.^92^ Decreased activity in frontoparietal regions correlates with poorer working memory.^93^ Frontocentral and default mode areas were most affected in our work. The observed correlation with deficits in overall cognitive abilities (MoCA, EpiTrack) is thus highly plausible, especially considering the crucial role of these large-scale networks in multiple forms of complex cognition, such as working memory, reasoning, and language processing.^94–96^ That is, a relatively lower Hurst exponent is associated with poorer cognitive performance due to the propagation of dysregulated excitatory activity, which manifests in noisier, less efficient processing.^97^ We also found a positive relationship between the progressive changes in the MoCA score and the Hurst exponent in TLE, supporting the notion of progressive cognitive decline over time.^49, 98^ Collectively, these findings suggest that the Hurst exponent might be a clinically useful index for monitoring the progression of cortical E/I imbalance and cognitive impairment in pharmaco-resistant TLE.

## Methods

### Participants

#### Discovery Dataset (MICA-MICs)

We studied 40 individuals with pharmaco-resistant TLE [17 males; mean ± SD age = 35.80 ± 11.04 years (18-63 years)], who underwent MRI examination for research purposes at the Montreal Neurological Institute and Hospital between 2018 and 2023. TLE diagnosis and lateralization of seizure focus as left (*n* = 27) and right TLE (*n* = 13) followed ILAE criteria,^55^ and were determined by a comprehensive evaluation that included detailed history, review of medical records, neuropsychological assessment, video-EEG recordings, and clinical MRI. The control group included 40 healthy adults with no history of neurological or psychiatric conditions [19 males; 34.25 ± 3.98 years (28 - 44 years)] who underwent MRI scans using the same imaging protocol as the TLE group.^54^ There were no differences in age (*t* = 0.84, *P* = 0.406) and sex (*χ*^2^ = 0.20, *P* = 0.653) between TLE and control groups. Detailed demographic and clinical information are provided in **Table 1**.

#### Replication Dataset (EpiC)

This dataset consisted of 30 pharmaco-resistant TLE patients [10 males; 30.87 ± 11.46 years (18 - 58 years)] who had undergone research-dedicated MRI scans at Universidad Nacional Autónoma de México between 2013 and 2017.^48, 51^ A similar evaluation classified patients as left (*n* = 11) or right TLE (*n* = 19). Patients were compared to 30 healthy adults [11 males; 31.83 ± 11.35 years (18 - 57 years)] who had no history of neurological or psychiatric illness and underwent the same imaging protocol. As in the discovery dataset, there were no differences in age (*t* = 0.33, *P* = 0.744) and sex (*χ*^2^ = 0.07, *P* = 0.787) between TLE and control groups. Detailed demographic and clinical information are provided in **Table 1**.

#### Standard Protocol Approvals and Patient Consents

Written informed consent was obtained from all participants. All studies were approved by local Ethics Review Committees and carried out according to the declaration of Helsinki.

### MRI Acquisition

#### Discovery Dataset (MICA-MICs)

All participants (healthy controls and patients) underwent baseline multimodal MRI scans, including T1-weighted, diffusion-weighted, and resting-state fMRI. Twelve patients underwent 1.98 ± 1.26 years of follow-up scans, of which five additionally underwent 3.40 ± 0.55 years of follow-up scans. All scans were acquired on a 3.0 T Siemens Magnetom Prisma-Fit scanner equipped with a 64-channel head coil. Two T1-weighted scans were acquired using a 3D MPRAGE sequence (TR = 2300 ms, TE = 3.14 ms, FA = 9°, FOV = 256×256 mm^2^, voxel size = 0.8×0.8×0.8 mm^3^, matrix size = 320×320, 224 slices). Diffusion-weighted MRI data were acquired using a 2D EPI sequence (TR = 3500 ms, TE = 64.4 ms, FA = 90°, FOV = 224×224 mm^2^, voxel size = 1.6×1.6×1.6 mm^3^, 3 b0 images, b-values = 300/700/2000 s/mm^2^ with 10/40/90 diffusion directions). Resting-state fMRI data were acquired using a multiband accelerated 2D EPI sequence (TR = 600 ms, TE = 30 ms, FA = 52°, FOV = 240×240 mm^2^, voxel size = 3×3×3 mm^3^, matrix size = 80 ×80, multi-band factor = 6, 48 slices, 700 volumes). A subset participant (28 controls and 27 patients) additionally underwent pseudo-continuous arterial spin labeling (ASL) MRI (TR = 4150 ms, TE = 10 ms, FA = 90°, voxel size = 4.5×4.5×7 mm^3^, FOV = 288×288 mm^2^, post-label delay = 1550 ms, 14 slices).

#### Replication dataset (EpiC)

All participants had multimodal MRI scans (T1-weighted, diffusion MRI and resting-state fMRI), of which 14 TLE patients had follow-up scans with an mean interscan interval of 2.61 ± 0.89 years (range = 0.7-4 years). All scans were acquired using a 3T Philips Achieva MR scanner, and included (i) one T1-weighted MRI scan (3D gradient-echo EPI, TR = 8.1 ms, TE = 3.7 ms, FA = 8°, FOV = 256×256 mm^2^, voxel size = 1×1×1 mm^3^, 240 slices), (ii) one resting-state fMRI scan (2D gradient-echo EPI, TR = 2000ms, TE = 30 ms, FA = 90°, voxel size = 2×2×3 mm^3^, 34 slices, 200 volumes), and (iii) one diffusion-weighted MRI scan (2D EPI, TR = 11.86 s, TE = 64.3 ms, FOV = 256×256 mm^2^, voxel size = 2×2×2 mm^3^, 2 b0 images, b-value = 2000 s/mm^2^, 60 diffusion directions).

### MRI Preprocessing

MRI data from the *MICA-MICs* and *EpiC* datasets were processed using virtually identical pipelines via *micapipe* (version 0.2.2; http://micapipe.readthedocs.io),^99^ an open multimodal MRI pipeline that integrates AFNI, FSL, FreeSurfer, ANTs, MRtrix, and Workbench.^100–104^ T1-weighted data underwent gradient non-uniformity correction, re-orientation, skull stripping, intensity normalization, and tissue segmentation. Diffusion-weighted data underwent denoising, b0 intensity normalization, and correction for susceptibility distortion, head motion, and eddy current. Resting-state fMRI processing involved discarding the first five volumes, re-orientation, motion, and distortion correction. Nuisance variable signals were removed using a ICA-FIX classifier.^105^ Volumetric time series were non-linearly co-registered to native FreeSurfer space with boundary-based registration,^106^ and mapped to individual mid-thickness surfaces with trilinear interpolation. Cortical time series were resampled to the Conte69 surface space (with ∼32k vertices/hemisphere) and smoothed with a 10-mm full-width half-maximum (FWHM) kernel. ASL MRI data were processed using FSL-BASIL (https://asl-docs.readthedocs.io). We used *oxford_asl*, an automated command line utility within BASIL, to generate a calibrated map of absolute resting-state tissue perfusion for each participant. For further details, see ref.^107^ Resulting cortical blood flow (CBF) map was co-registered to the native FreeSurfer space using boundary-based registration,^106^ projected onto the Conte69 surface space, and smoothed using a 10-mm FWHM kernel. Lastly, subject-specific vertex-wise resting-state fMRI time series and CBF maps were parcellated into 360 cortical regions defined by the HCP multi-modal parcellation (HCP-MMP).^108^

### Connectivity Matrix Generation

Functional connectivity (FC) was calculated as the correlation coefficient of the fully processed time series for each pair of regions (360×360). A functional connectivity dynamics (FCD) index was computed as follows. Each region’s time series (with a total length of 695 time points) were segmented into 596 windows of 100 time points each (60 s), with an overlap of 99 time points. The whole-brain FC matrix was constructed for each time window and vectorized by considering only the upper triangular entries. These vectorized matrices were then cross-correlated, generating a 596×596 FCD matrix for each participant.

Individual structural connectivity (SC) was generated from preprocessed diffusion-weighted data via MRtrix.^104^ Anatomically-constrained tractography was first performed using tissue types (cortical and subcortical gray matter, white matter, cerebrospinal fluid) derived from each participant’s processed T1-weighted images registered to native DWI space.^109^ Multi-shell and multi-tissue response functions were estimated, and constrained spherical deconvolution and intensity normalization were performed.^110, 111^ The tractogram was then generated based on a probabilistic approach with 40 million streamlines, with a maximum tract length of 250 and a fractional anisotropy cut-off of 0.06. Subsequently, spherical deconvolution-informed filtering of tractograms (SIFT2)^112^ was applied to reconstruct the whole-brain streamlines weighted by cross-sectional multipliers. Lastly, the SC matrix (360×360) was constructed by mapping the reconstructed cross-section streamlines onto the HCP-MMP atlas with 360 nodes, in which the connection weights between nodes were defined as the weighted streamline count.

### Hurst Exponent Analysis

Prior work using *in-silico* modeling and *in-vivo* chemogenetic manipulations has validated the utility of the Hurst exponent, an index estimated from neural time-series data,^5^ to infer the underlying change in the synaptic E/I ratio. In these reports, the Hurst exponent decreased as the E/I ratio shifted toward higher excitation. Here, we calculated the Hurst exponent of each brain region’s preprocessed restingstate fMRI time series and used it as a proxy of the overall E/I ratio within that area. In brief, for each participant, each brain region’s time series were modeled as fractionally integrated processes, and the corresponding Hurst exponent was estimated using the univariate maximum likelihood method and discrete wavelet transform.^5,113^ The specific function utilized is the *bfn_mfin_ml.m* function from the *nonfractal* MATLAB toolbox (https://github.com/wonsang/nonfractal),^114^ with the ‘filter’ argument set to ‘haar’ and the ‘ub’ and ‘lb’ arguments set to [1.5, 10] and [-0.5, 0], respectively.

To contextualise our regional pattern of the Hurst exponent to a range of molecular, structural, and functional features, we obtained relevant brain maps from the literature using BigBrainWarp toolbox (https://bigbrainwarp.readthedocs.io)^115^ and neuromaps (https://netneurolab.github.io/neuromaps/)^59^. We fetched and parcellated the map of the sensory-fugal axis of cytoarchitectural differentiation,^116^ cortical thickness,^72^ intracortical myelination (T1w/T2w ratio),^117^ gene expression gradient,^60^ and neurotransmitter receptor gradient.^61, 69^ Spearman rank correlations separately quantified the relationship between each brain annotation and the group-level Hurst exponent map in healthy controls. Statistical significance of spatial correlation between brain maps (*i.e.*, *P*_spin_) was assessed non-parametrically via comparison against a null distribution of null maps with preserved spatial autocorrelation,^66^ *i.e.*, spin tests with 5,000 iterations, implemented using the ENIGMA Toolbox (https://enigma-toolbox.readthedocs.io).^67^

Before statistical analysis, region-wise Hurst exponent values in TLE patients were initially normalized relative to healthy controls, and sorted into ipsilateral/contralateral to the epileptogenic focus.^118^ Surface-based linear models assessed group differences in each brain area’s Hurst exponent between patients and healthy controls using BrainStat (https://brainstat.readthedocs.io).^119^ Effect size was calculated as Cohen’s *d*. We controlled for age and sex, and we corrected findings for false discovery rate (FDR) using random field theory for nonisotropic images.^120, 121^ For regions surviving FDR correction, we conducted *post-hoc* analyses using two-sample *t*-tests. To examine network-wise group differences, the 360 brain regions of the whole-brain were grouped into twelve macroscale networks according to the Cole-Anticevic brain-wide network partition (CAB-NP) defined on the HCP-MMP atlas.^62^ We then calculated the average Hurst exponent for each network of each individual, and compared them between groups using two-sample *t*-tests. To demonstrate the robustness of our findings with respect to head motion, we calculated the mean framewise displacement (FD) from resting-state fMRI scans for each participant. We further repeated the surface-wide comparison of the Hurst exponent while additionally controlling for the mean FD.

### Recurrent Neural Circuit Modeling

A biophysically-based mean-field model was used to simulate coordinated neuronal activities across the whole brain based on long-range anatomical connection and to infer microcircuit-level parameters of neuronal populations at a regional level. Specifically, we harnessed a parametric mean-field model (pMFM)^26^ that captures the link between time-varying functional dynamics of intrinsic brain activity and structural connection, as well as its modulation through region-specific microcircuit parameters. In comparison to other models that also incorporate local microcircuit properties,^64, 122^ the pMFM, by allowing them to vary along the anatomical and functional hierarchical axes of the cerebral cortex,^26, 60^ generate more realistic simulations of large-scale brain dynamics with modest parametric complexity. A comprehensive description of pMFM, including the mathematical details of the mean-field model, can be found in refs.^26, 64, 123^ In brief, the pMFM assumes that the neural dynamics of a given brain region are governed by four components: (i) recurrent (intra-regional) input, where a larger recurrent input current corresponds to a stronger recurrent connection strength *w*; (ii) inter-regional input, mediated by structural connection strength between a pair of regions and scaled by a global scaling constant *G*; (iii) external input *I*, mainly from subcortical structures; (iv) neuronal noise, assumed to be Gaussian with a standard deviation *σ*. Here, recurrent connection strength *w*, external input current *I*, and noise amplitude *σ* varied across brain regions, while *G* was kept a constant. Additionally, *w*, *I*, and *σ* were parameterized as linear combinations of T1w/T2w myelin maps^117^ and the principal gradient of resting-state functional connectivity,^124^ rather than varying independently:

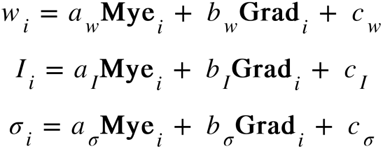

where *w_i_*, *I_i_*, and *σ_i_* were the recurrent connection strength, external input current, and noise amplitude of the *i*-th cortical region, respectively. Mye*_i_* and Grad*_i_* denoted the average scores of T1w/T2w MRI estimates of intracortical myelin and the principal resting-state FC gradient in the *i*-th cortical region. Therefore, there are a total of 10 unknown parameters (*a_w_*, *b_w_*, *c_w_*, *a_I_*, *b_I_*, *c_I_*, *a_σ_*, *b_σ_*, *c_σ_*, *G*) to be estimated by maximizing fit to empirical static FC and FCD (**Fig. 2a**).

In this study, for each group (healthy controls, or TLE), 40 participants were randomly subdivided into training (*n* = 15), validation (*n* = 15), and test (*n* = 10) sets. Group-level SC and FC matrices (360×360) were computed by averaging the FC and SC matrices across participants separately within the training, validation, and test sets. FCD matrices could not be directly averaged across participants as there was no temporal correspondence of resting-state time series between subjects. The cumulative distribution function (CDF) of every participant’s FCD matrix was constructed by collapsing the upper triangular entries, and then simply averaged across all participants separately within the training, validation, and test sets, which we referred to as a group FCD CDF.^26^ Subsequently, in the training set, the CMA-ES algorithm^65^ was iterated 50 times, and repeated 5 times with different random initializations, yielding 250 candidate parameter sets. The 250 candidate parameter sets were evaluated in the validation set. The top 10 candidate parameter sets selected from the validation set based on the model fit were tested in the test set to determine the optimal set of parameters (with the lowest cost). More specifically, the simulated fMRI signal from each parameter set was used to compute a 360×360 static FC matrix and a 596×596 FCD matrix. The agreement between the simulated and empirical static FC matrices was defined as Pearson’s correlation (*r*) between the upper triangular entries of the two matrices, in which a larger *r* indicated a more similar static FC. The disagreement between the simulated and empirical FCD matrices was defined as the Kolmogorov–Smirnov (KS) distance^125^ between the two matrices’ CDF, in which a smaller KS distance indicated a more similar FCD. Following previous work, an overall cost was defined as [(1 − *r*) + KS] to optimize both static FC and FCD;^26^ lower cost thus implied a better fit to empirical static FC and FCD. For robustness, the split of participants into training, validation, and test sets was repeated five times for each group. Lastly, the best parameter sets from these five splits were averaged, yielding the representative set of parameters for each group.

Spearman correlations were calculated between the group-level pFMF parameters (**Fig. 2b**) and Hurst exponent (**Fig. 1a**) to evaluate the association between regional Hurst exponent changes and microcircuit properties. Regional changes in microcircuit parameters (*w* and *I*) between TLE and controls were quantified by simply subtracting their group-level parameter scores. These changes were correlated with the effect size of regional differences in the Hurst exponent (*i.e.*, Cohen’s *d* in **Fig. 1b**). Significances of spatial correlations were determined via spin permutations tests, with 5,000 iterations.

### Network Spreading Mapping

Group-average SC and FC matrices derived from an independent sample (*i.e.*, HCP) of 100 unrelated healthy participants were used to estimate the mean abnormality of neighbours of each brain region.^72^ Briefly, neighbours of a given brain region *i* were defined as regions connected to it with a structural connection, as defined by the SC matrix. The structurally-connected neighbour abnormality of node *i* (*D_i_*) was estimated as the average weighted abnormality of all of *i*’s all neighbours,^69^ where *d_j_* is the abnormality (*i.e.*, Cohen’s *d* score) of the *j*-th neighbour of node *i*, *SC_ij_* is the SC strength between node *i* and node *j*, and *N_i_* is the total number of neighbours that are connected to node *i* (*i.e.*, node degree).

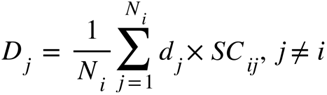

Structurallyand functionally-defined neighbour abnormality was estimated using the same equation as above, with the exception that regional abnormality was additionally weighted by the FC strength to node *i* (*FC_ij_*):^69^

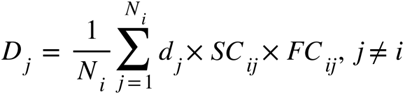

Altogether, a single neighbour abnormality value was estimated for each condition of each brain area. Spearman correlation coefficients were used to assess the relationship between node abnormality and mean abnormality of structurally-defined neighbours, and both structurally- and functionally-defined neighbours, separately. Spatial autocorrelation-preserving spin tests were used to assess the statistical significance of associations across brain regions.

### Disease Epicenters Mapping

Disease epicenters were identified by spatially correlating each brain region’s healthy structural and functional connectivity profiles (from the same HCP dataset) to our map of Hurst exponent alterations in TLE (*i.e.*, un-thresholded Cohen’s *d* map in **Fig. 1b**).^71^ This approach was repeated systematically across all cortical regions with spin permutation tests at *P*_spin_ < 0.05. The higher the spatial similarity between a node’s connectivity profile and the whole-brain patterns of Hurst exponent disruption, the more likely this structure represented a disease epicenter (**Fig. 3b**), regardless of its abnormality level. Resulting likelihoods (*i.e.*, correlation coefficients) were then ranked in descending order, with highly ranked brain regions representing disease epicenters.

### Associations with Clinical and Cognitive Variables

For those regions showing significant between-group differences (**Fig. 1b**), we assessed the effect of disease severity (age at seizure onset, disease duration, and number of antiepileptic drugs) on the level of Hurst exponent changes in TLE patients. The analyses were performed in the cross-sectional and longitudinal cohorts combined (57 MRI scans in total). We fitted linear mixed-effects models containing *participant* intercept as a random term, and each clinical variable as a fixed term,^43^ and tested for a negative effect of the given clinical variable. Associations between the Hurst exponent and the number of electroclinical seizures captured during hospitalization were examined using Pearson’s *r*. In addition, we explored the effect of interictal epileptiform discharges (IEDs) in the temporal lobe on Hurst exponent changes in patients who underwent extended video-EEG telemetry (mean ± SD = 8.68 ± 2.58 days, range = 2-15 days). For each TLE patient, we obtained the IEDs prevalence based on the classification from clinical EEG reports during hospitalization. Following the ACNS Critical Care EEG Terminology 2021,^126^ TLE patients were divided into 2 subgroups: rare IEDs (<1/hour, *n* = 11) and prevalent IEDs (*i.e.*, occasional/frequent/abundant, ≥1/hour, *n* = 27). Differences in the Hurst exponent between the two subgroups were tested using two-sample *t*-tests.

To unveil the neurophysiological substrate of Hurst exponent changes observed, we conducted further analysis on cortical blood flow (CBF). CBF is tightly linked to brain metabolism,^127, 128^ varies across the lifespan,^129, 130^ and is increasingly recognized as a key neuroimaging biomarker for various neuropsychiatric and neurological disorders.^127, 131^ In this work, 28 healthy controls and 27 patients underwent ASL MRI. To compare regional CBF with corresponding Hurst exponent values, we computed each region’s average CBF score and Hurst exponent in healthy controls, then calculated Spearman’s correlation between them; the *P*-value was determined via spin permutation tests with 5,000 iterations. Additionally, we assessed how well TLE-related changes in the Hurst exponent reflected CBF changes. We separately computed the global average CBF and the average CBF in brain regions showing significant Hurst exponent changes (**Fig. 1b**) for each participant. Subsequently, we (1) compared CBF values between TLE patients and healthy controls using two-sample *t*-tests, and (2) correlated CBF with subject-specific Hurst exponent values while controlling for age and sex.

Neuropsychological assessments at the time of study MRI were also available for most participants, including general cognitive function [Montreal Cognitive Assessment (MoCA);^74^ 39 controls and 38 patients], and attention and executive functions (EpiTrack;^75^ 38 controls and 28 patients). We directly compared TLE patients to healthy controls using two-sample *t*-tests. To examine the clinical significance of cortical E/I imbalance, we separately extracted the Hurst exponent for the entire brain and in those significant areas, then correlated them with the individual cognitive measurements described above, controlling for age and sex. In a separate analysis of TLE patients for whole longitudinal data was available, we quantified progressive changes in Hurst exponent, MoCA, and EpiTrack scores by calculating the differences between baseline and follow-up pairs, and examined their associations.

### Replication Analysis

Hurst exponent alterations between TLE patients and healthy controls were validated in an independent dataset (*i.e.*, *EpiC*; 30 controls and 30 TLE) to verify the robustness of our findings. The protocols used for the analyses were the same as those described above. Briefly, we separately compared differences in the average Hurst exponent across the entire brain and only within the significant regions identified in the discovery sample (see **Fig. 1b**) between TLE patients and healthy controls using twosample *t*-tests. Surface-wide differences in the Hurst exponent between TLE and control groups were assessed via surface-based linear models. The spatial correspondence between *MICA-MICs* and *EpiC* datasets for the effect sizes (*i.e.*, Cohen’s *d*) of Hurst exponent changes was examined with spin permutation tests (5,000 iterations). We proceeded to investigate the effect of disease severity on Hurst exponent changes across time. We separately calculated the average Hurst exponent across the entire brain or within significant regions from the follow-up scans, for each patient for whole longitudinal data was available (*n* = 14), then compared them to those derived from the baseline scans. We also assessed the relationship between the Hurst exponent and epilepsy duration in TLE across both crosssectional and longitudinal scans using linear mixed-effects models that contained *participant* intercept as a random term and *epilepsy duration* as a fixed term.

Finally, we validated brain-behavior associations. At the *EpiC* site, a subset of participants (21 controls and 28 patients) underwent the Wechsler Memory Scale (WMS-IV) test that consisted of 7 subtests designed to assess memory performance.^51^ Every subject’s performance is reported as 5 index scores: auditory memory (AMI), visual memory (VMI), visual working memory (VWMI), immediate memory (IMI), and delayed memory (DMI). These indices were normalized with respect to a Mexican population and adjusted for age and education level, then were entered into a principal component analysis to reduce dimensionality. The loading score of the first component (*i.e.*, PC1), accounting for 81% of the variance, was correlated with individual Hurst exponent values, with age and sex as covariates.

## Data Availability

The *MICA-MICs* dataset^54^ is available at the Open Science Framework (https://osf.io/j532r/) and the Canadian Open Neuroscience Platform (https://portal.conp.ca/). The *EpiC* dataset is openly available at OpenNeuro (data set ds004469, https://openneuro.org/datasets/ds004469/versions/1.1.3). Surfacebased Hurst exponent data of the study participants will be available on osf.io upon publication.

## Code Availability

MRI preprocessing was conducted using micapipe (http://micapipe.readthedocs.io).^99^ The Hurst exponent was computed using the nonfractal toolbox (https://github.com/wonsang/nonfractal). Surfacebased statistics were conducted using BrainStat (https://brainstat.readthedocs.io).^119^ Spin permutation tests were conducted using the ENIGMA Toolbox (https://enigma-toolbox.readthedocs.io).^67^ Computational modeling was conducted using pMFM (https://github.com/HeavenBluer/Parametric-MFM-Project).^26^

## Acknowledgements

K.X. is funded by the China Scholarship Council (CSC). J.R. is funded by the Canadian Institutes of Health Research (CIHR). R.R.C., A.N., E.S., and J.C. are funded by the Fonds de Recherche du Québec - Santé (FRQ-S). L.C. is funded by the CONACYT (181508, 1782) and UNAM-DGAPA (IB201712, IG200117, IN204720). B.C.B. acknowledges research support from the National Science and Engineering Research Council of Canada (NSERC Discovery-1304413), CIHR (FDN-154298, PJT-174995), SickKids Foundation (NI17-039), Helmholtz International BigBrain Analytics and Learning Laboratory (HIBALL), Healthy Brains and Healthy Lives (HBHL), Brain Canada, FRQS, and the Tier-2 Canada Research Chairs program.

## Supplementary Information

**Fig. S1.**
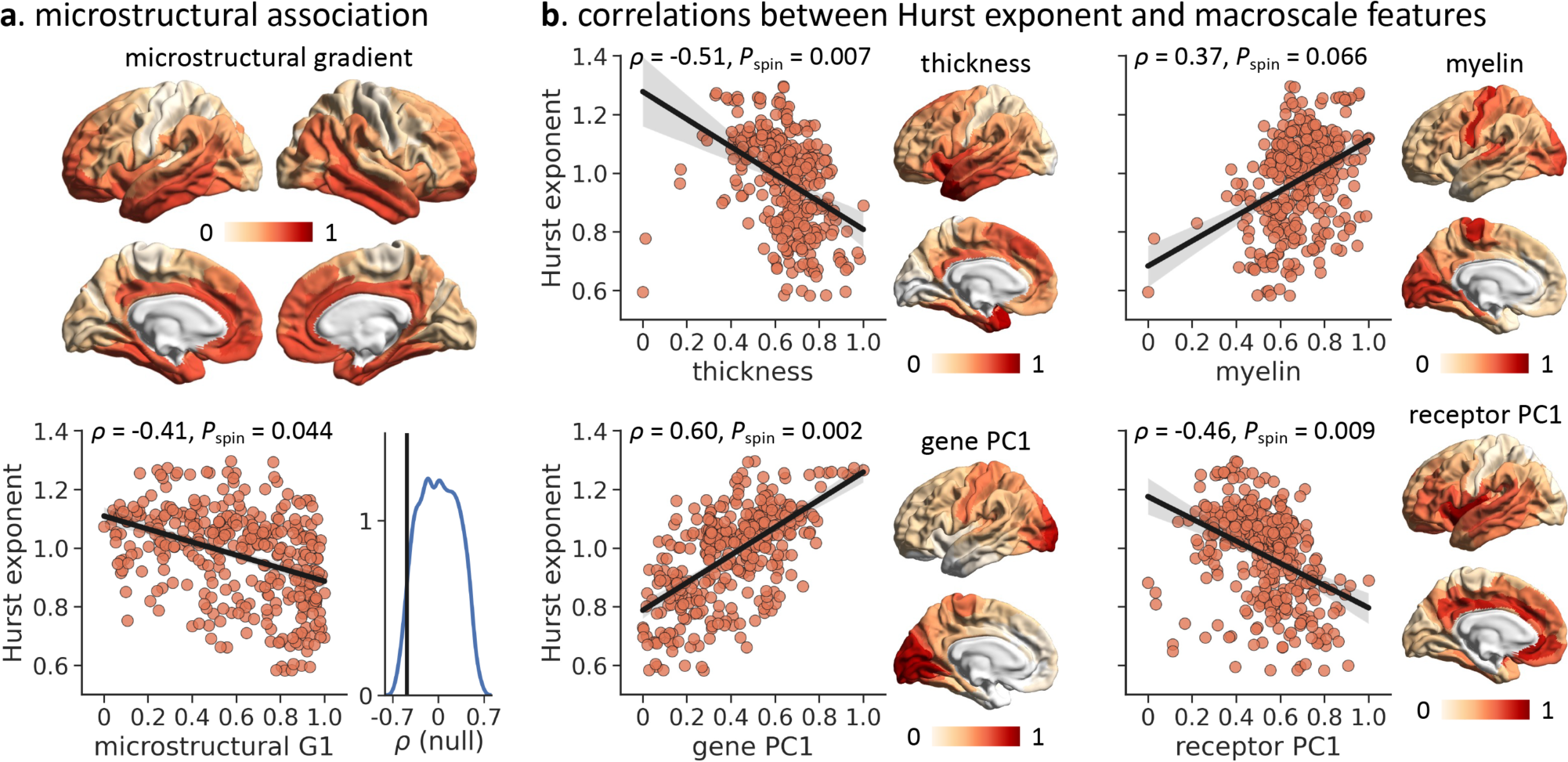
Regional Hurst exponent values vary spatially across the brain. **(a)** Regional Hurst exponent value is negatively spatially correlated with the sensory-fugal hierarchy of cytoarchitectural differentiation (“BigBrain gradient”). **(b)** Regional Hurst exponent value spatially aligns with the regional distribution of cortical thickness, intracortical myelination (T1w/T2w MRI ratio), the principal component of AHBA brain-specific gene expression data (“gene PC1”), and the principal component of the density of neurotransmitter transporters/receptor (“receptor PC1”). For example, a lower Hurst exponent (*i.e.*, greater E/I ratio) is found in brain regions with lower intracortical myelination. The statistical significance of spatial correlation between brain maps (*i.e.*, *P*_spin_) was assessed non-parametrically using spin permutation tests (5,000 iterations) that preserved spatial autocorrelation.

**Fig. S2.**
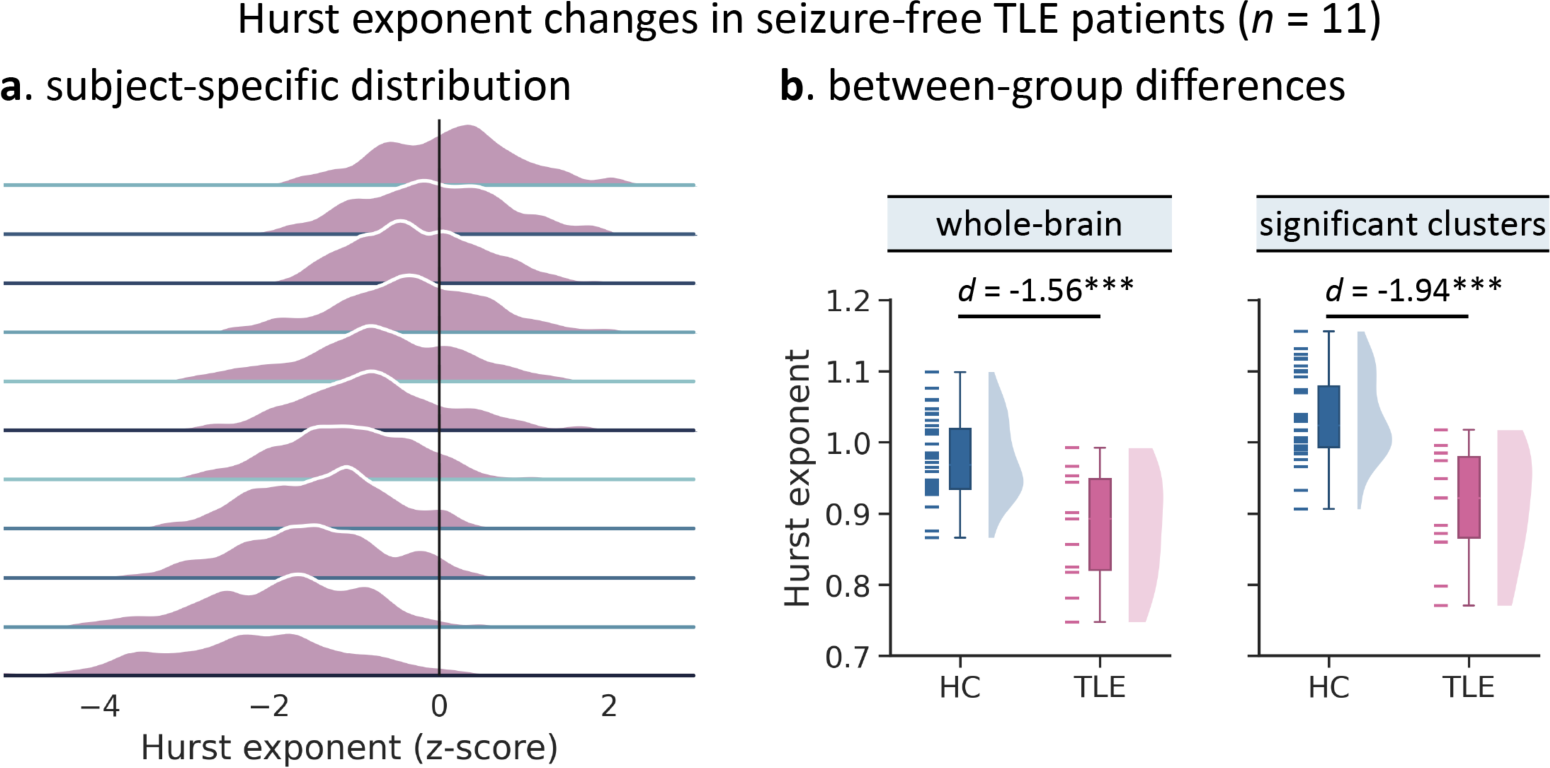
Hurst exponent changes in only seizure-free TLE patients. **(a)** Subject-specific distribution of regional Hurst exponent (*z*-score relative to healthy controls) in seizure-free TLE patients (*n* = 11). **(b)** Between-group differences in the average Hurst exponent values across the entire brain (*left*) or significant brain regions (*right*) in Fig. 1b. *** *P* < 0.001. HC = healthy control; TLE = temporal lobe epilepsy.

**Fig. S3.**
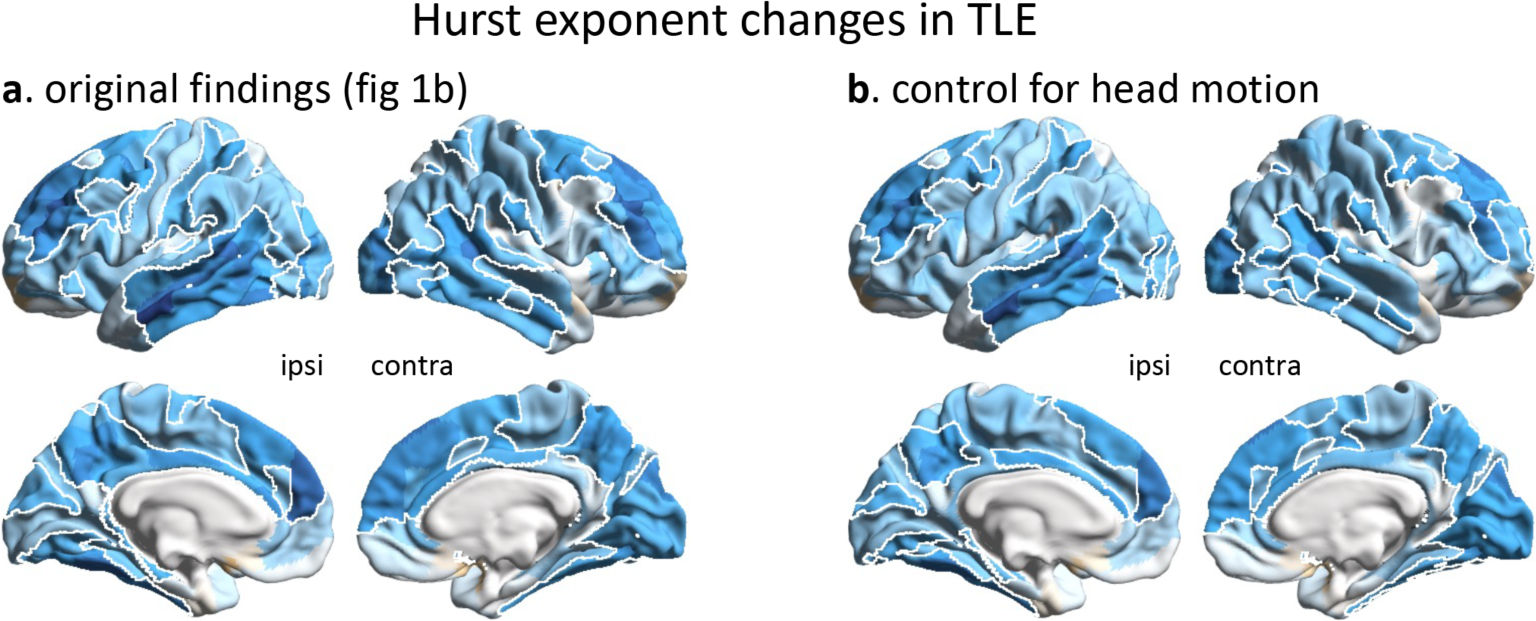
Head motion effects. Hurst exponent differences between TLE patients and healthy controls with (**b**) or without (**a**) additionally controlling for individual head motion. Surface-based findings were corrected for multiple comparisons at a false discovery rate level of 0.05 (*P*_FDR_ < 0.05; white outlines). ipsi = ipsilateral; contra = contralateral.

**Fig. S4.**
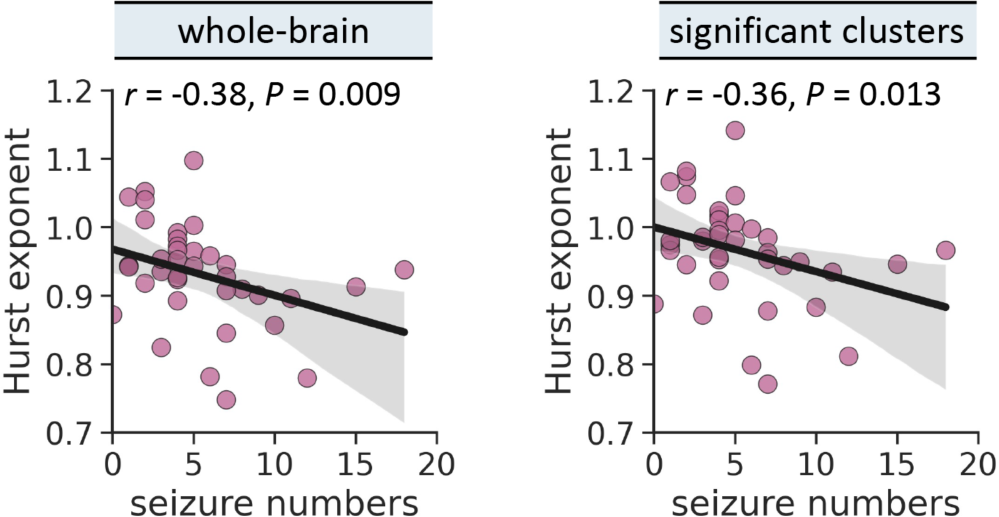
Correlations between the average Hurst exponent and the number of electroclinical seizures captured during the EMU admission in TLE patients. Subject-specific mean Hurst exponent value is calculated by averaging the Hurst exponent values across the entire brain (*left*) or significant brain regions (*right*) in Fig. 1b.

**Fig. S5.**
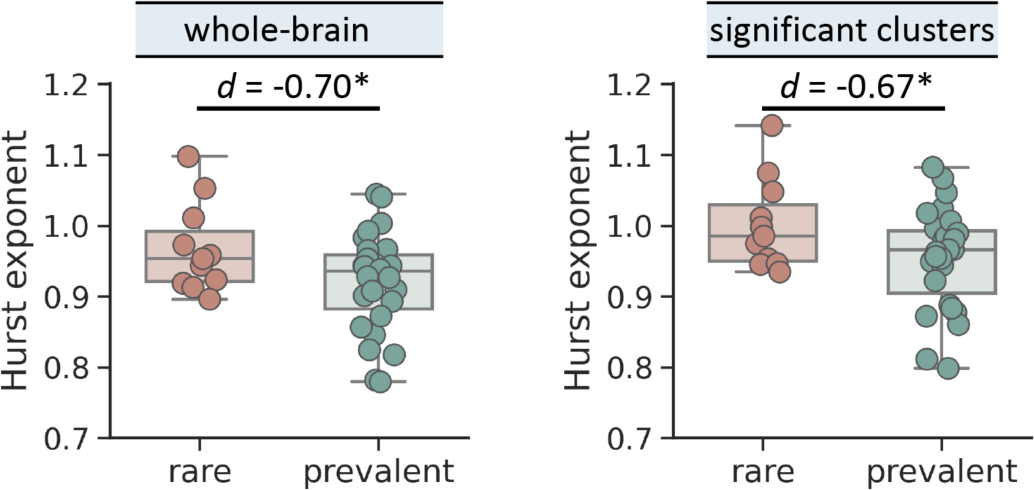
Interictal epileptiform discharges (IEDs) effects. Differences in the average Hurst exponent between patients with rare (*n* = 11) and prevalent (*i.e.*, occasional/frequent/abundant, *n* = 27) interictal epileptic discharges (IEDs). Subject-specific mean Hurst exponent value is calculated by averaging the Hurst exponent values across the entire brain (*left*) or significant brain regions (*right*) in Fig. 1b.

**Fig. S6.**
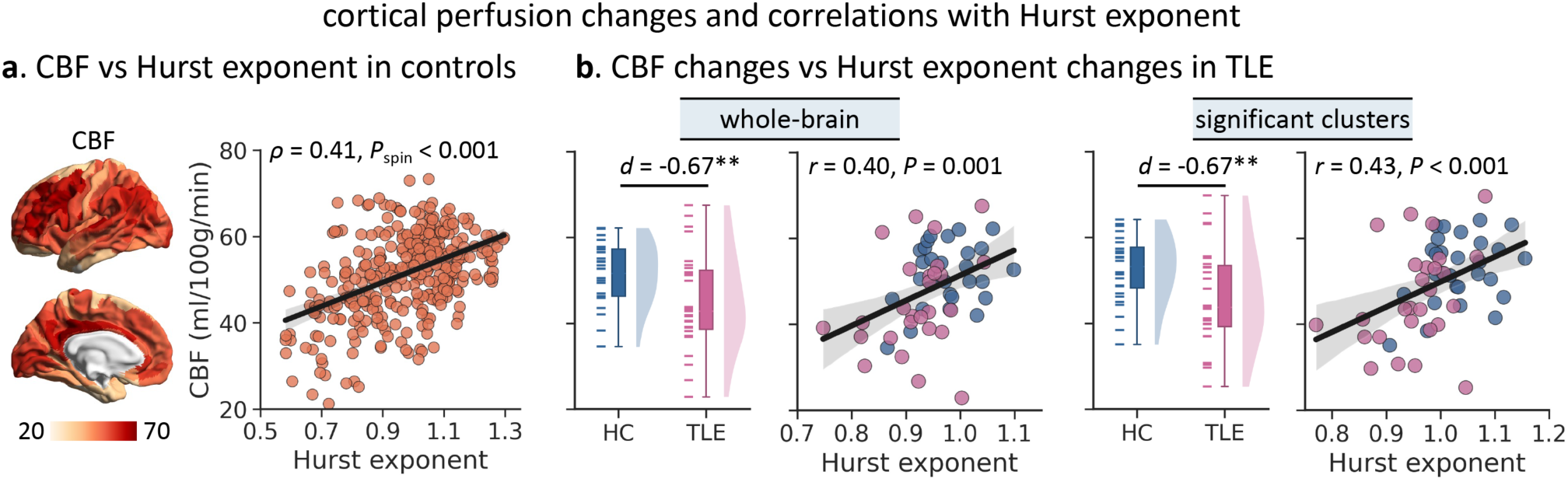
Regional brain perfusion changes in TLE and association with Hurst exponent changes. **(a)** Regional Hurst exponent value spatially aligns with the spatial distribution of regional cortical blood flow (CBF) derived from restingstate arterial spin labeling (ASL) MRI. A lower Hurst exponent (*i.e*., greater E/I ratio) is seen in brain regions with lower brain perfusion. The statistical significance of the spatial correlation between brain maps (*i.e.*, *P*_spin_) was assessed non-parametrically by spin permutation tests (5,000 iterations) that preserved spatial autocorrelation. **(b)** Error bar plots: between-group differences in the average CBF scores across the entire brain (*left*) or brain regions (*right*) showing significant Hurst exponent changes (Fig. 1b). Scatter plots: associations between subject-specific average Hurst exponent and CBF. ** *P* < 0.010. HC = healthy control; TLE = temporal lobe epilepsy.

